# Mycobacterial methionine aminopeptidase type 1c moonlights as an anti-association factor on the 30S ribosomal subunit

**DOI:** 10.1101/2024.11.26.625572

**Authors:** Aneek Banerjee, Krishnamoorthi Srinivasan, Jayati Sengupta

## Abstract

Methionine aminopeptidase (MetAP), an essential metalloprotease, binds the 70S ribosome near the exit site of peptide tunnel and cleaves the N-terminal methionine from the nascent polypeptide chains. While *E. coli* expresses a single subclass of type1 MetAP, mycobacteria possesses two subclasses, namely, MetAP1a and MetAP1c, differentiated by the presence of a 40 amino acid long N-terminal tether in the later. Here we reveal a yet unknown moonlighting role of MetAP1c in mycobacteria. We observe stationary phase-specific enhancement of expression and a unique 30S subunit affinity of MetAP1c. A 5Å cryo-EM map of *Mycobacterium smegmatis* (Msm) MetAP1c-30S subunit complex manifests for the first time the binding of MetAP1c at the inter-subunit side in the vicinity of proteins S13 and S19. MetAP1c binding and resulting conformational changes make the 30S subunit unfit for association with the 50S subunit, thereby conferring an anti-association property on MetAP1c. We further constructed two N-terminal mutants to establish the pivotal role of the N-terminal extension of MetAP1c in its moonlighting function.

## Introduction

Methionine aminopeptidase (MetAP), exist in all kingdoms of life (1), is a divalent cation-based metalloprotease. It is the key player for N-terminal methionine excision (NME) where it cleaves the initiator methionine from almost 70% of nascent polypeptide chains thus generating diversity in the N-terminal peptide (2) (3). Structurally, MetAP family proteins contain a conserved ‘pita-bread’ fold (4) (Fig S1A). Depending on the absence or presence of a 60 amino acid insertion in the catalytic core, MetAP is divided into type I and type II respectively (5). Type I MetAP is prevalent in both prokaryotes (subclasses Ia, Ic) and eukaryotes (subclasses Ib, Id), whereas type II is found only in eukaryotes (3) (6) (Fig S1B).

Interestingly, while most of the prokaryotes contain only MetAP1a gene, actinomycetes have a second MetAP1c gene. MetAP1a and MetAP1c in mycobacteria are differentiated on the basis of absence or presence of a 40 amino acid N-terminal extension (7) (Fig S1C). As type 1c is unique to mycobacteria, it poses as an attractive species-specific drug target. Small molecules and drugs targeting mycobacterial MetAP1c have been recently identified (8) (9). Various biochemical studies highlighted that both MetAP1a and MetAP1c are enzymatically active (10) (11). The *in-vitro* enzymatic activity varies depending on the type of substrate as both the proteins display subtle substrate preference (12). Interestingly, the N-terminal extension of MetAP1c plays a crucial role in maintaining its methionine excision property. Although the extension is not a part of the catalytic core, removal of the 40 amino acid extension abolishes the enzymatic activity of MetAP1c (12). Deletion of the gene coding for MetAP1a results in slow growing mycobacteria but deleting the gene encoding MetAP1c protein results in decreased survivability of mycobacteria which signifies the importance of type 1c protein (13). Despite some progress in understanding, the exact reason for maintaining the two subclasses of MetAP in mycobacteria is still elusive. Moreover, the importance of MetAP1c in mycobacteria as observed raised the question of whether this protein could have additional functions beyond N-terminal methionine excision.

To better understand the functional role of MetAPs, we have analyzed the growth phase-dependent expression profile of the two subclasses of *M. smegmatis* (Msm) MetAP’s and their binding preferences towards the ribosomal subunits. Surprisingly, our results revealed that MsmMetAP1c expression increases in late stages of growth phase and it showed remarkable affinity towards the 30S ribosomal subunit. We obtained a cryo-EM map of MsmMetAP1c-30S subunit complex at 5Å resolution showing an extra density, attributable to MetAP1c, protruded at the inter-subunit face. Taking a cue from this first structure of MsmMetAP1c-30S complex, we demonstrated a moonlighting anti-association function of MetAP1c on the 30S subunit. Constructing two N-terminal mutants, we further identified a pivotal role of the 40 residues N-terminal extension present in MetAP1c in mediating the anti-association activity.

Overall, here we present a novel mechanism of how MsmMetAP1c binds to the 30S subunit *via* its N-terminal extension and marks the 30S-MetAP1c complex unfavorable for associating with the 50S subunit to form active 70S ribosome, and thereby down regulates the rate of protein translation. We propose, this could be a unique mechanism by which mycobacteria sequester 30S subunit during stationary phase of growth or under stress conditions to regulate the rate of translation.

## Results

### Mycobacterial MetAP’s show preferential binding to the ribosomal subunits

Structurally MetAP1a and MetAP1c are very similar with a conserved catalytic core allowing them to function as methionine excision machinery. The major difference between mycobacterial MetAP1a and MetAP1c is the presence of a 40 amino acid long N-terminal extension in MetAP1c (Fig.1A, B). We checked the expression pattern of these two proteins in various growth phases of *Mycobacterium smegmatis* (Msm). We analyzed the protein levels in early log, late log and stationary phases of bacterial growth using anti-MetAP1a and anti-MetAP1c antibodies. In line with the observation on mRNA levels of the proteins in *M. tuberculosis* (10), which showed that the mRNA level gets elevated during log phase for MetAP1a while for MetAP1c during stationary phase of growth, MsmMetAP1a level is maximum during log phase of growth and it decreases as the cell grows older (Fig.1C, D). The result is reverse in case of MsmMetAP1c where low protein level during early log phase and highest during stationary phase was observed (Fig.1E, F). This phase-specific inverse relation between MsmMetAP1a and MsmMetAP1c expression further stroked our interest and prompted us to check the affinity of the proteins for ribosome and its subunits.

**Figure 1.**
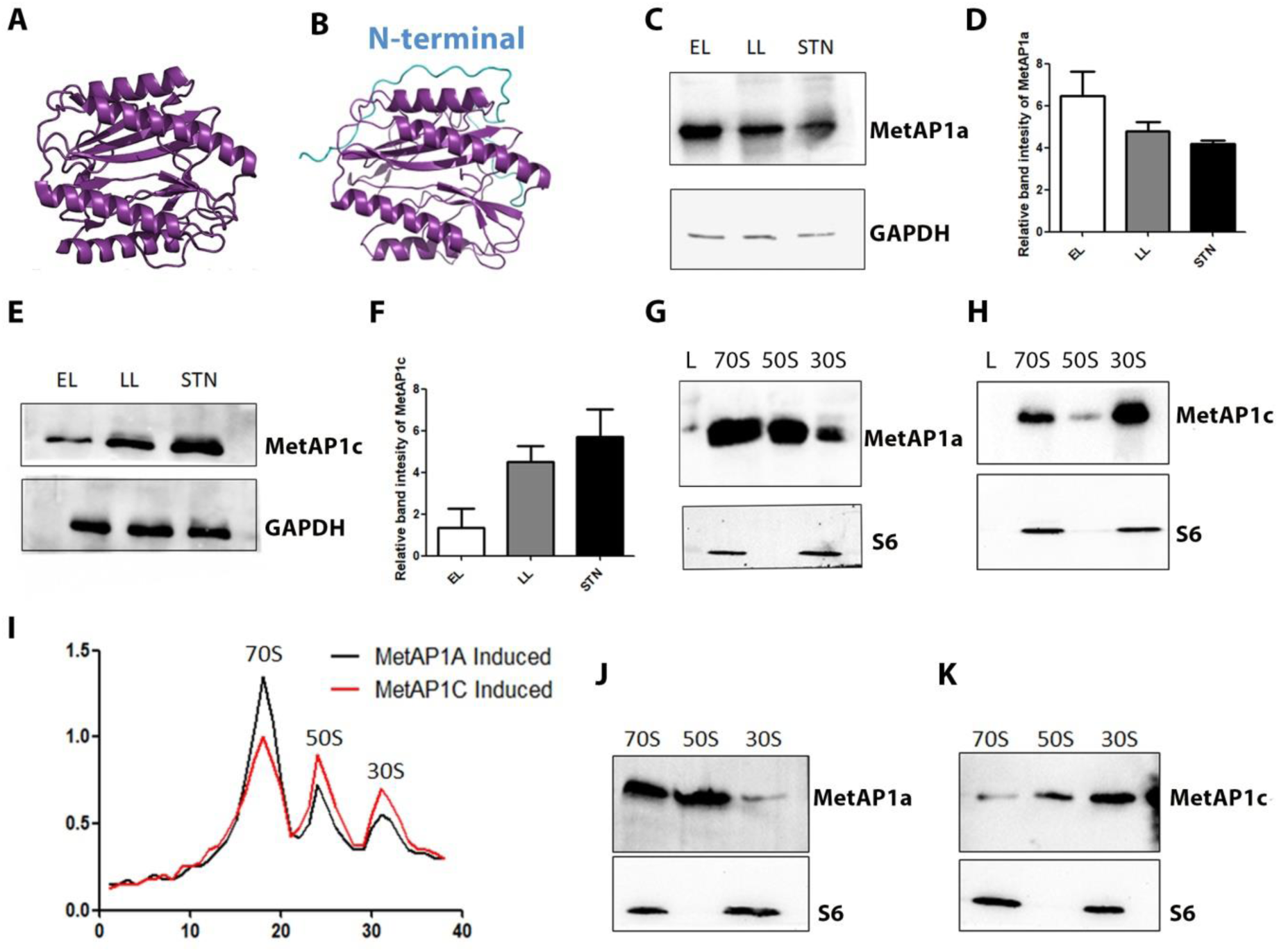
Growth-phase specific expression and preferential binding of mycobacterial MetAP1c to ribosome and ribosomal subunits. The Phyre2 predicted structures of MsmMetAP1a (A) and MsmMetAP1c (B) emphasize the presence of the N-terminal extension (marked in blue) unique to type 1c methionine aminopeptidase. The phase-specific expression of MsmMetAP1a is determined *via* western blot analysis (C), with GAPDH serving as the loading control. Changes in level of expression are depicted in the bar plot (D). Similarly, the expression of MsmMetAP1c is assessed by western blot (E) and represented in a bar diagram (F). The *in-vitro* ribosomal subunit preference of MsmMetAP1a and MsmMetAP1c is illustrated in panels (G) and (H), respectively. Small ribosomal subunit protein S6 was used as a loading control for ribosome. The ribosomal profile following induction of MsmMetAP1a and MsmMetAP1c in *M. smegmatis* cells (I) reveals a decrease in the 70S population after MsmMetAP1c induction. Fraction analysis via western blotting (J) mimics the *in-vitro* observations, with MsmMetAP1a displaying preferential binding to the 70S ribosome and 50S subunit, while MsmMetAP1c favors binding to the 30S subunit (K). (In the ribosomal profile of the panel I, X axis corresponds to fraction number and Y axis corresponds to OD260)

Methionine aminopeptidases are known to bind near the exit site of the peptide tunnel on the 50S subunit. Accordingly, MsmMetAP’s are expected to show higher affinity to the 70S ribosome as well as to the 50S subunit and for MsmMetAP1a this was the observed binding pattern as seen in the western blot analysis (Fig.1G). MsmMetAP1c, in contrast, showed higher binding affinity to 30S subunit (Fig.1H). To check if the same profile of binding preference exists *in-vivo*, we cloned both the genes into pLam12 vector, expressed them in *M. smegmatis* cells and purified the crude ribosome (Fig.1I). Analyzing the ribosomal fractions revealed a similar result with MsmMetAP1a preferring the large ribosomal subunit and MsmMetAP1c favoring the small subunit (Fig.1J, K). Of note, one interesting result that caught our attention was the lowering of 70S population upon MetAP1c induction (Fig. 1I). Adding that to the protein’s increased affinity to the 30S subunit strongly hinted towards its anti-association role on the 30S subunit. Similar *in-vivo* assay was done using *E. coli* cells expressing mycobacterial MetAP’s and it demonstrated identical result (Fig.SD-F). The increased expression of MsmMetAP1c during stationary phase of growth and its binding preference to the 30S ribosomal subunit intrigued us to determine its binding location on the 30S subunit.

### Cryo-EM structure of the MsmMetAP1c-30S subunit complex

Cleaving the initiator methionine from the nascent polypeptide is the only known function of bacterial methionine aminopeptidase and to perform that, the protein needs to access the peptide exit tunnel of the 70S ribosome on the 50S subunit. However, biochemical assays manifested higher affinity of MsmMetAP1c towards 30S subunit indicating a hitherto unknown moonlighting function of the protein which we wanted to explore. To that end, we solved a cryo-EM structure of MsmMetAP1c-30S subunit complex at ∼5Å resolution (Fig.2A, S2A, B and Table S1). An additional density, attributable to MetAP1c, was visible at the inter-subunit side surface of the 30S subunit (Fig. 2A). The molecular model of the 30S subunit with the additional protein density has been built. Crystal structure of *M. tuberculosis* MetAP1c is available and MsmMetAP1c model was built based on that as they share a sequence similarity of over 90% (Fig S2C). The 30S subunit head region is known to be inherently dynamic (14), which was notably amplified in the presence of MsmMetAP1c, as evidenced by the 2D and 3D class analyses (Fig S3). The ligand density appeared less resolved. The reason could be that the protein with conserved pita-bread fold has a propensity to bind structured RNAs (15) (16). Thus its interaction with the rRNA generates conformational heterogeneity in head region of the 30S subunit. It appeared that the interactions with the ribosomal proteins eventually stabilize MetAP1c on the inter-subunit surface of the 30S subunit to some extent.

**Figure 2.**
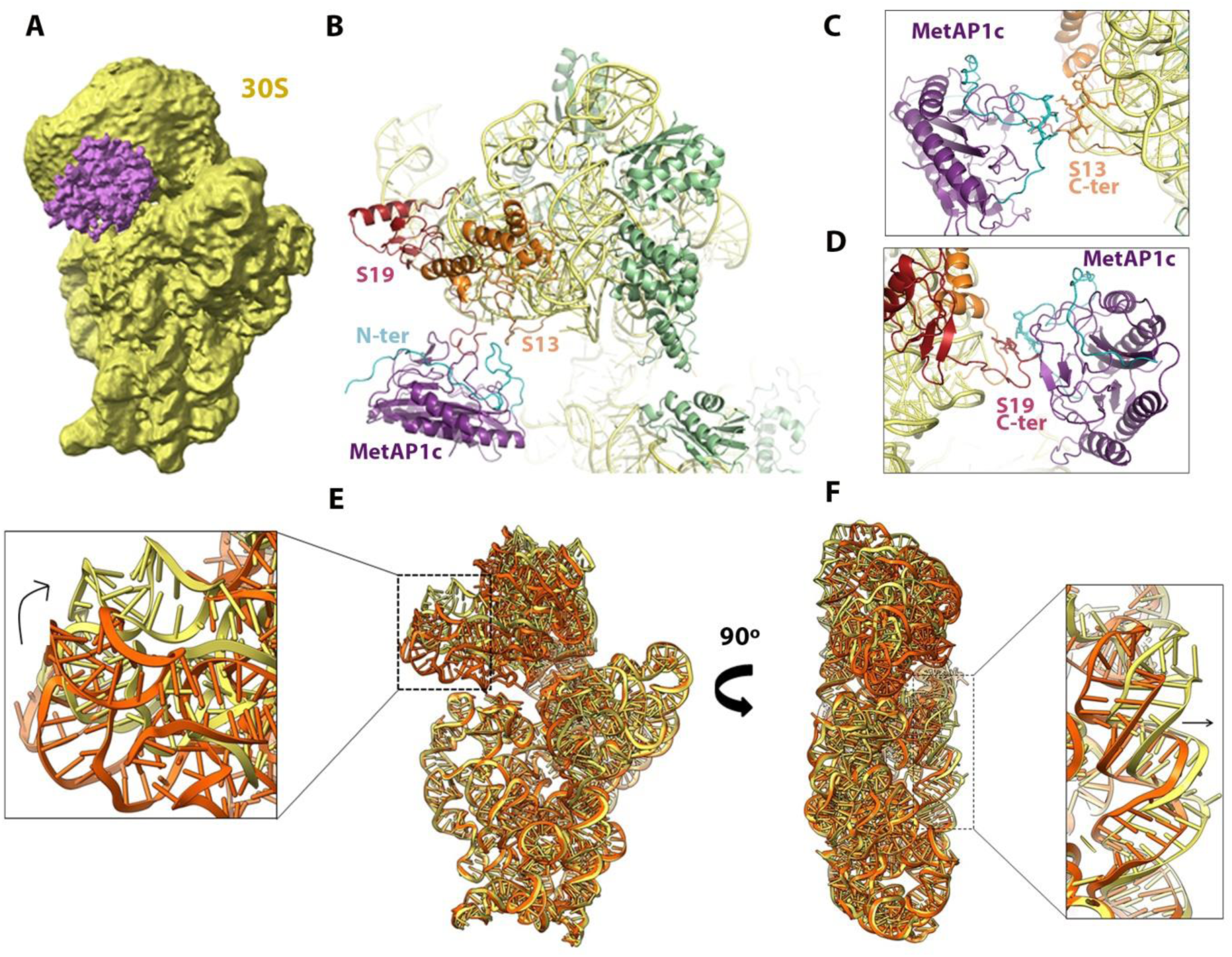
Cryo-EM structure of the MsmMetAP1c-30S subunit complex. MetAP1c (colored in purple) is visualized in the 5Å cryo-EM map of the MsmMetAP1c-30S subunit complex. The 30S subunit is shown in yellow (A). Cartoon representation of the molecular model of MsmMetAP1c binding site accentuates the close proximity of MsmMetAP1c (purple) to ribosomal proteins S13 (orange) and S19 (brick-red) (B). Close-up views of the interaction region reveal potential interaction of N-terminal extension (cyan) of MsmMetAP1c with the C-terminus of S13 (C) and S19 (D). Comparison between the 16S rRNA core of a ligand-free 30S subunit (PDB: 5ZEU, orange) and our model (yellow) highlights a head tilt of approximately 11° (E) and about 10Å displacement of the helix 44 near the decoding center (F).

**Figure 3.**
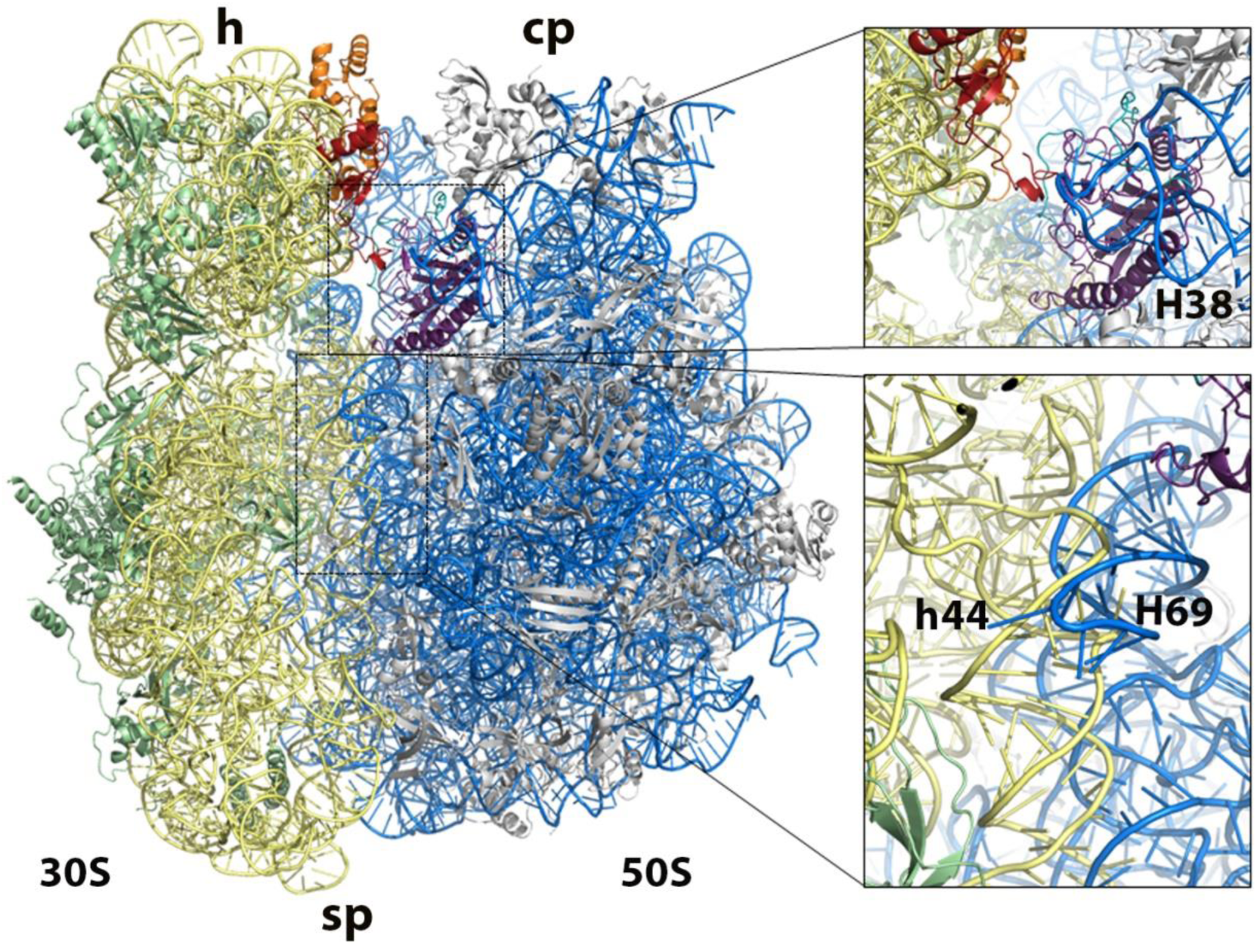
Binding of MsmMetAP1c makes the 30S subunit structurally incompatible for 70S ribosome formation. Alignment of the structure of MsmMetAP1c-30S subunit complex with a 70S ribosome (PDB code: 5O61) reveals potential impediments to 70S formation efficiency (left panel). In the upper right panel, close up view clearly shows a steric clash between MsmMetAP1c and the 50S subunit (shown in blue). Binding of MetAP1c alters the conformation of helix 44 (protruded towards inter-subunit side), leading to unfavorable contacts with Helix 69 of the 23S rRNA, preventing the bridge formation between the two subunits, as depicted in the lower right panel. The 16S rRNA is shown in yellow; S13 in orange, S19 in brick red and all other small subunit proteins are in green; 23S and 5S rRNA is in blue; all large subunit proteins are in grey. MsmMetAP1c is in purple. Landmarks: h, Head; sp, Spur; cp, Central Protuberance

MetAP1c anchors onto the inter-subunit face of the 30S subunit head, in the immediate neighborhood of the 16S rRNA helix 44 (h44), through its N-terminal extension and interacts with ribosomal proteins S13 and S19 (Fig.2B). Apparently, amino acids 30 to 35 of MsmMetAP1c N-terminal extension are in close proximity to the C-terminal of ribosomal protein S13 (Fig.2C). The C-terminal tail of S19 also lies in the vicinity of the N-terminal region of MetAP1c (Fig.2D). The binding of MetAP1c alters the conformation of 16S rRNA of the 30S subunit.

During protein synthesis, the 30S subunit associates with the 50S subunit to form active 70S ribosome (17). The conformation of the head region of 30S subunit changes when it forms the 70S ribosome (18). Comparing the 16S rRNA model of a 30S subunit isolated from a ligand free 70S ribosome (PDB 5ZEU, (19)) with our structure, highlighted two major conformational differences: i) The beak region of the 30S subunit head is rotated by approximately 11° resulting in overall tilting of the head (Fig.2E). ii) Helix44 of the 16S rRNA is protruded out significantly by about 10Å towards the inter-subunit region (Fig.2F). Helix 44 (h44) is a crucial rRNA segment which forms major inter-subunit bridges (B2a, B3, B5) by interacting with the 23S rRNA helices of the large subunit (20). These bridges are crucial to the formation and to maintain stability of active 70S ribosome.

Alignment with a 70S ribosome structure (PDB: 5O61, (21)) demonstrated that the position of MsmMetAP1c binding would result in steric clash with the 50S subunit and the protrusion of h44 would make the inter-subunit bridge formation with the Helix69 of 23S rRNA unfavorable (Fig3). This observation piqued our interest to inspect whether MsmMetAP1c does act as an anti-association factor or not by binding to the 30S subunit.

### MsmMetAP1c moonlights as an anti-association factor on the 30S subunit

Our structural data strongly suggested that MsmMetAP1c-bound 30S subunit would not be able to associate with the 50S subunit to form active 70S ribosome. To test that hypothesis, we pre-incubated Msm30S subunit with MetAP1c and added equimolar Msm50S subunit for association (Fig.4A). The mixture was loaded on top of sucrose density gradient and ultra-centrifuged to obtain ribosomal fractions. Indeed, the profile clearly indicated that with increase in concentration of MsmMetAP1c there is a decrease in the 70S ribosome formation (Fig.4B-D). Light scattering assay results also supported this observation. With the addition of 10 times and 15 times MsmMetAP1c the amplitude of scattering decreased which hinted towards smaller particle size due to lower association of the 30S and 50S subunits (Fig.4E). To further validate the anti-association property of MsmMetAP1c, we performed the sucrose density gradient ultracentrifugation with the *E. coli* 30S and 50S subunits. It was observed that MetAP1c is even capable of prohibiting *E. coli* ribosomal subunits to join pointing towards a universal mechanism of anti-association activity of MsmMetAP1c (Fig.S4A-C). Notably, our observation of *in-vivo* MsmMetAP1a and MsmMetAP1c preferential binding to the ribosomal subunits, previously shown (Fig. 1), is in agreement with the above findings. The reason behind lowering of 70S population upon MsmMetAP1c induction is indeed due to the anti-association effect of the protein. Over-expression of MsmMetAP1c leads to its enhanced binding to the 30S subunit making the complex unfit for associating with the 50S subunit. To check whether this anti-association is a unique property of MsmMetAP1c, we performed similar sucrose density gradient-based subunits association studies with MsmMetAP1a and *E. coli* MetAP (EcMetAP). The obtained ribosomal profile, in contrast, indicated the absence of any anti-association effect of both of these proteins (Fig.4F-K, S4D-F).

**Figure 4.**
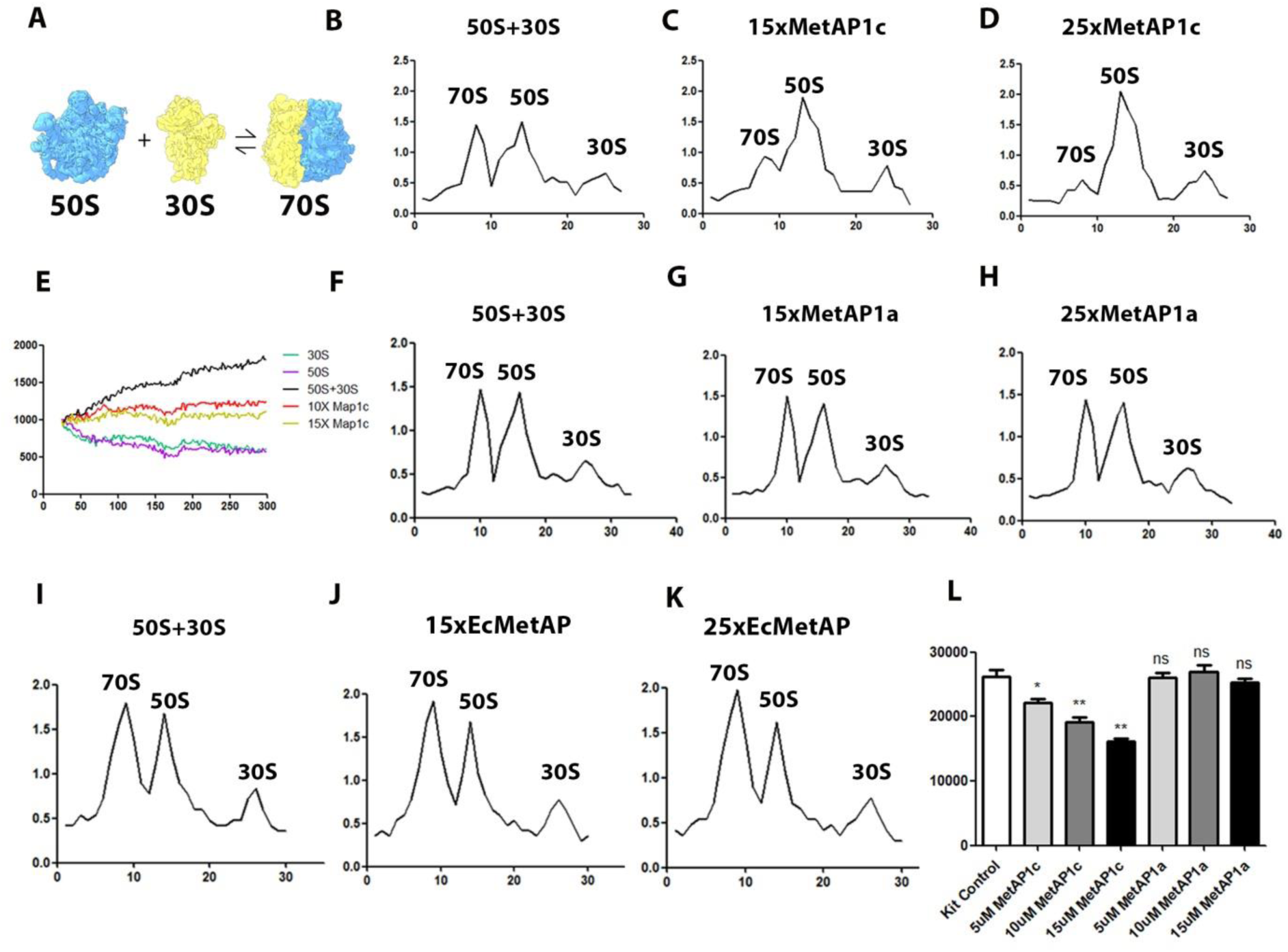
MsmMetAP1c binding to the 30S subunit inhibits the formation of 70S ribosome. A schematic representation of the small subunit and the large subunit association to form 70S ribosome (A). The formation of 70S ribosome, as evidenced by sucrose density gradient profile in (B), occurs through the association of 50S and 30S subunits. (C) and (D) illustrate the impact of adding 15-fold and 25-fold excess MsmMetAP1c to the 30S subunit, respectively, resulting in reduced 70S ribosome formation. This observation is supported by the light scattering assay in (E), where increase in protein concentration leads to decrease in scattering intensity. Association assays using MsmMetAP1a reveal comparable ribosomal profiles between the control set (F) and those with 15-fold (G) and 25-fold (H) excess MsmMetAP1a, suggesting that the subunit association is not inhibited by the presence of the protein. Similar results are observed with *E. coli* MetAP (EcMetAP). The association profile of the control set (I) closely resemble those with 15-fold and 25-fold excess EcMetAP (J, K) respectively. The amount of luciferase synthesis is analyzed by measuring the firefly luminescence (L). High concentration of MsmMetAP1c results in lower luminescence indicating reduced translation due to lower 70S formation, whereas MsmMetAP1a has no effect. (X axis corresponds to fraction number and Y axis corresponds to OD260 in the association profiles)

Prevention of 70S formation should down-regulate the rate of translation. Indeed, the change in the level of luciferase synthesis by adding increasing amount of MsmMetAP1c showed that the elevation in MsmMetAP1c concentration led to a reduction in the amount of luciferase synthesis, while MsmMetAP1a had no discernible effect on luciferase levels (Fig.4L). Clearly, anti-association role by binding to the 30S ribosomal subunit is a moonlighting property of MsmMetAP1c, which EcMetAP as well as MsmMetAP1a lack. A paramount structural difference between these proteins is the presence of a 40 amino acid extension in MsmMetAP1c (6). Next, we checked whether this additional segment plays any role in anti-association.

### N-terminal extension of MsmMetAP1c plays a crucial role in its anti-association activity

Cryo-EM map of MsmMetAP1c-30S subunit complex presented here revealed that this N-terminal extension lies within the interaction zone of ribosomal proteins S13 and S19. To look into the role of the N-terminus of MsmMetAP1c in anti-association, we designed two mutants, one by deleting the 40 amino acid N-terminal extension of MsmMetAP1c (Del40 MetAP1c) and another one by adding the N-terminal extension upstream of MsmMetAP1a (Chimeric MetAP1a) (Fig.5A, B). The mutant proteins were purified to determine their anti-association activity. 30S subunits were pre-incubated with Del40 MetAP1c and Chimeric MetAP1a followed by addition of equimolar amount of 50S subunit for association. Ribosomal profile was obtained after sucrose density gradient ultracentrifugation of the reaction mixture. For Del40 MetAP1c, there was no change in 70S ribosome formation with the addition of the mutant protein to both mycobacterial and *E. coli* ribosomes (Fig.5C-E, S5A-C). The anti-association property was abolished in Del40 MetAP1c thus validating the significance of the N-terminal extension. Western blot of the ribosomal fractions from 25 times protein treated set highlighted the shift of binding preference from 30S subunit to 50S subunit and 70S ribosome (Fig.5F). Interestingly, the chimeric MetAP1a displayed a gain of anti-association activity as the amount of 70S ribosome declined with the increase in protein concentration (Fig.5G-I). The lowering of 70S formation was also prominent when *E. coli* subunits were used (Fig S5D, E). Furthermore, the binding affinity of this chimeric MetAP1a showed a significant enhancement towards the 30S ribosomal subunit (Fig.5J). We then analyzed the level of luciferase production while adding increasing amount of Del40 MetAP1c and chimeric MetAP1a. Going in-line with our previous observations, chimeric MetAP1a reduced the luciferase production while Del40 MetAP1c did not cause any effect (Fig S5F). Combing all the biochemical and structural data, it is quite evident that the 40 amino acid N-terminal extension of MsmMetAP1c contributes significantly towards the protein’s preferential binding to 30S subunit which in-turn prevents the association with 50S subunit to form active 70S ribosome.

**Figure 5.**
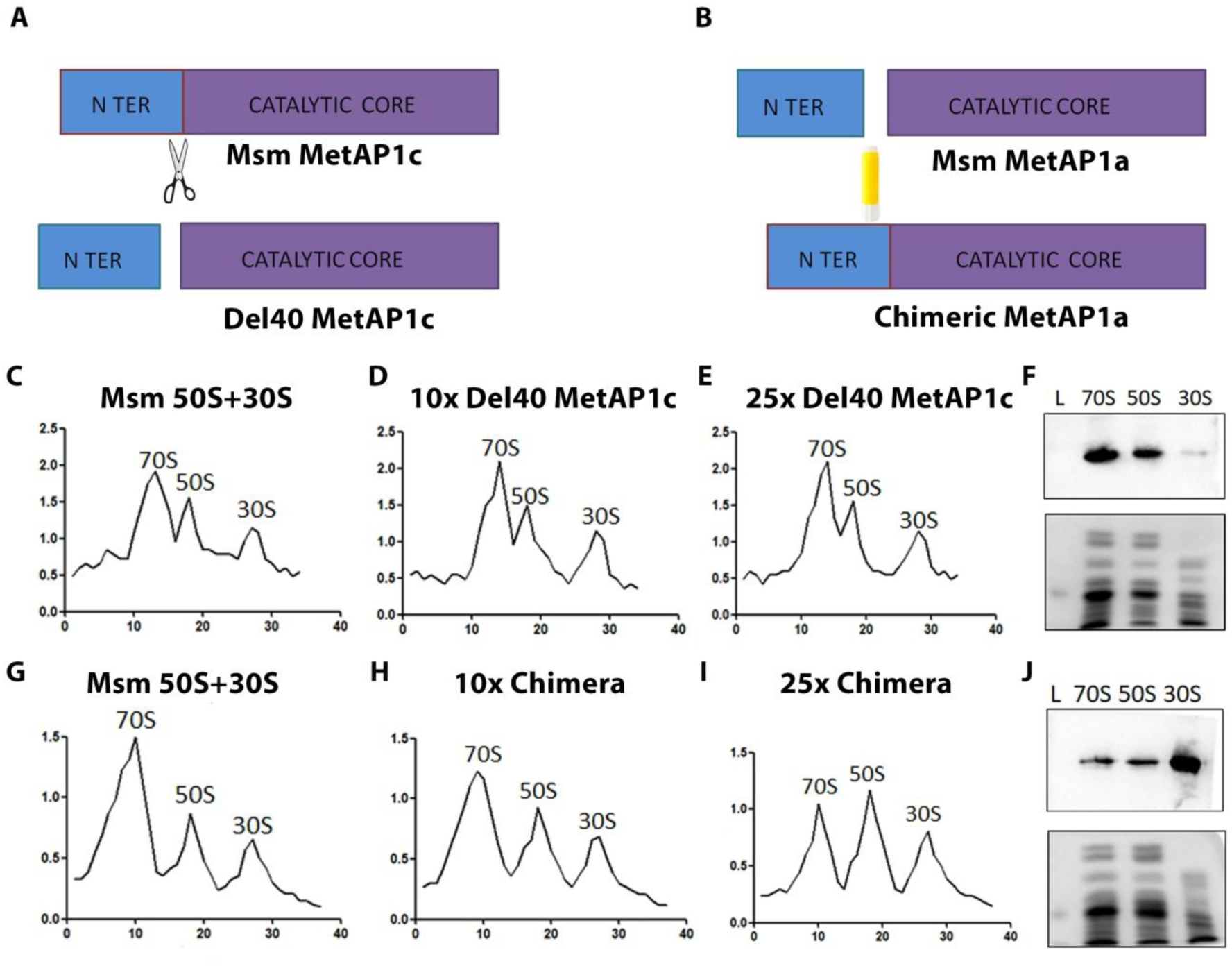
The 40 amino acid long N-terminal extension of MetAP1c is essential for its anti-association property. To validate the significance of the N-terminal extension, two mutant proteins were designed. Del40 MetAP1c (A) represents 40 amino acid N-terminal extension deleted form of MsmMetAP1c, while Chimeric MetAP1a (B) displays addition of the 40 amino acids upstream of MsmMetAP1a. Ribosomal subunit association assays were conducted using both mutant proteins. Compared to the control set (C), 10-fold (D) and 25-fold (E) excess Del40 MetAP1c shows no reduction in 70S formation. Western blot analysis of the 25-fold excess protein fractions reveals a shifted binding preference towards the 70S ribosome and 50S subunit (F). Chimeric MetAP1a exhibits a gain of anti-association property. Relative to the control (G), 10-fold (H) and 25-fold (I) excess Chimeric MetAP1a display decreased 70S ribosome formation from 50S and 30S subunits. Interestingly, Chimeric MetAP1a demonstrates higher binding affinity to the 30S subunit (J) when analyzed by western blotting. (In the association profiles, X axis corresponds to fraction number and Y axis corresponds to OD260)

## Discussion

The primary function of methionine aminopeptidase (MetAP), a divalent metal cation based co-translational metalloprotease, is to cleave the initiator methionine from roughly 50%-70% of the nascent polypeptides during translation (2) (3). As a co-translational nascent chain processing enzyme, MetAP needs to access the exit site of ribosomal peptide tunnel. A cryo-EM map of *E. coli* 70S ribosome bound to MetAP and peptide deformylase (PDF) highlighted how the proteins interplay among themselves to bind near the exit tunnel and perform their enzymatic function in *E. coli* (22) (23) . Interestingly, *Mycobacterium smegmatis* (Msm) possesses two subclasses of type 1 MetAP, type 1a containing just the catalytic core (similar to *E. coli* MetAP) whereas, type 1c containing a 40 amino acid long N-terminal extension in addition to the catalytic core (6). Both of these proteins were shown to be enzymatically active although having different substrate preference (12). The preferred substrate of MetAP1a was Met-Gly-Met-Met whereas for MetAP1c it was Met-Ala-Ser (12). We attempted to determine the structure of the MsmMetAP1c-70S complex in order to explore its potential involvement in co-translational events. However, our efforts did not yield any MetAP1c-bound complex. One possible explanation for this outcome could be the requirement for additional factors that facilitate the binding of MsmMetAP1c near the peptide tunnel exit, as observed in eukaryotic systems where the Nascent-polypeptide Associated Complex (NAC) serves to recruit MetAP1 by offering a platform for binding near the exit tunnel (24). However, our study unveiled a hitherto unknown function of MetAP1c in mycobacteria and provided insight into a mechanism of biological adaptation of mycobacteria in translational regulation.

There was a long-standing debate as to which subclass of MsmMetAP (MsmMetAP1a or MsmMetAP1c) is essential in mycobacteria, and only recently it has been established that MsmMetAP1c is indispensable for its survival (13). Nevertheless, the rationale behind maintaining two types of MetAPs remained elusive. We delved into this realm and to that end we first investigated the growth phase-specific expression of MsmMetAP1a and MsmMetAP1c. Our results revealed a notable enhancement in the expression of MsmMetAP1c during the stationary phase, aligning with previous findings suggesting phase-specific alterations in mRNA levels for both type of MetAP’s in *M. tuberculosis* (10). Recent studies have reported similar growth phase-specific variations in MetAP levels for the protozoan parasite *Eimeria tenella*, where MetAP1 was found to be up-regulated during the merozoite stage, playing a crucial role in host cell invasion (25).

Furthermore, a distinct specificity of MsmMetAP1c towards the 30S subunit as revealed in this study has not been reported previously. Intriguingly, induction of MsmMetAP1c led to a reduction in the population of 70S ribosomes in contrast to MsmMetAP1a induction. Given that bacteria encounter nutritional starvation, hypoxia and various other stresses during stationary phase (26), our results collectively suggest a potential moonlighting function of MsmMetAP1c on the 30S ribosomal subunit.

Apparently, the N-terminus of MsmMetAP1c assists to anchor on the 30S subunit through interactions with the ribosomal proteins S13 and S19 as visualized in the cryo-EM map of MsmMetAP1c-30S subunit complex presented here. MsmMetAP1c binding to the small ribosomal subunit not only arrests the 16S rRNA in an altered conformation and does not allow the 30S subunit conformational changes required to form the 70S ribosome, but also sterically hinder association with the 50S subunit.

MetAP1c binding site is close to that observed in case of mycobacterial-specific protein Y (Mpy) which plays a role in ribosomal hibernation during zinc depletion (27). MetAP2 binding to the small subunit (near the protein S12) of the 80S ribosome has been shown recently (28). Interestingly, MetAP2 binds to the 40S subunit only after auto-proteolysis of its N-terminus. However, in contrast to the binding of MetAP1c, Mpy or MetAP2 binding does not hamper the 70S/80S ribosome formation or stability (Fig. S6A, B). In addition, the position of MsmMetAP1c on the 30S subunit has the capacity to cause steric clash with translation initiation factors and f-Met tRNA (29). Biochemical results, presented here, displaying decreased amount of 70S ribosome formation in the presence of increasing amount of MsmMetAP1c also hints towards an anti-association role of the protein.

Anti-association, a phenomenon that prevents the inter-subunit bridge formation between the 30S and 50S subunit, hinders active 70S ribosome formation. MsmMetAP1c hampers the crucial inter-subunit bridge formation and thus disrupts the association of ribosomal subunits. No other MetAP’s are reported to act as an anti-association agent which we validated using MsmMetAP1a and *E. coli* MetAP. The major difference between MsmMetAP1c with MsmMetAP1a and EcMetAP is the presence of a 40 amino acid N-terminal extension of MsmMetAP1c. The cryo-EM map presented here also suggested the involvement of this extension in 30S subunit anchoring. In agreement with our structural observation, it was observed that upon removal of these 40 amino acid residues (Del40 MetAP1c) MetAP1c loses its binding affinity to the 30S subunit and lacks anti-association activity. On the other hand, a mutant MsmMetAP1a, expressing the 40 amino acid N-terminal extension upstream of its catalytic core (Chimeric MetAP1a), displayed anti-association activity with enhanced binding to the 30S subunit. All these biochemical data, supported by our cryo-EM map, confirmed that the unique anti-association property of MsmMetAP1c is mediated by its N-terminal 40 amino acid extension.

Proteins of methionine aminopeptidase family and also proteins resembling similar folds as of MetAP, like EBP1, are known to play diverse roles in cellular processes (30). MetAP2 is a crucial regulator of cell proliferation and involved in the development of several cancers (31). Vasculogenic mimicry (VM), formation of new blood vessels around cancer cells, confers aggression and metastatic property to tumors. MetAP2 is also reported to play a significant role in this process and drugs are being identified to prevent VM (32). Taking together, it is clear that MetAPs are involved in a diverse range of biological processes which are slowly coming to light.

In mycobacteria, MetAP1c is up-regulated during stationary phase of growth. This phase mimics unfavorable growth condition and the bacteria adapt to adjust to environmental stresses (33). MetAP1c, functioning as an anti-association factor, can prevent active 70S ribosome formation and thus regulate the rate of translation (Fig.6). In *E. coli*, hibernation promoting factor (HPF) interacts with ribosome to form hibernating 100S ribosomes that are incapable of protein synthesis (34) (35). Mycobacteria lack this 100S formation mechanism (36) and it is conceivable that these anti-association factors like MetAP1c help the organism to regulate the rate of protein synthesis thus changing the dynamics from growth to survival.

**Figure 6.**
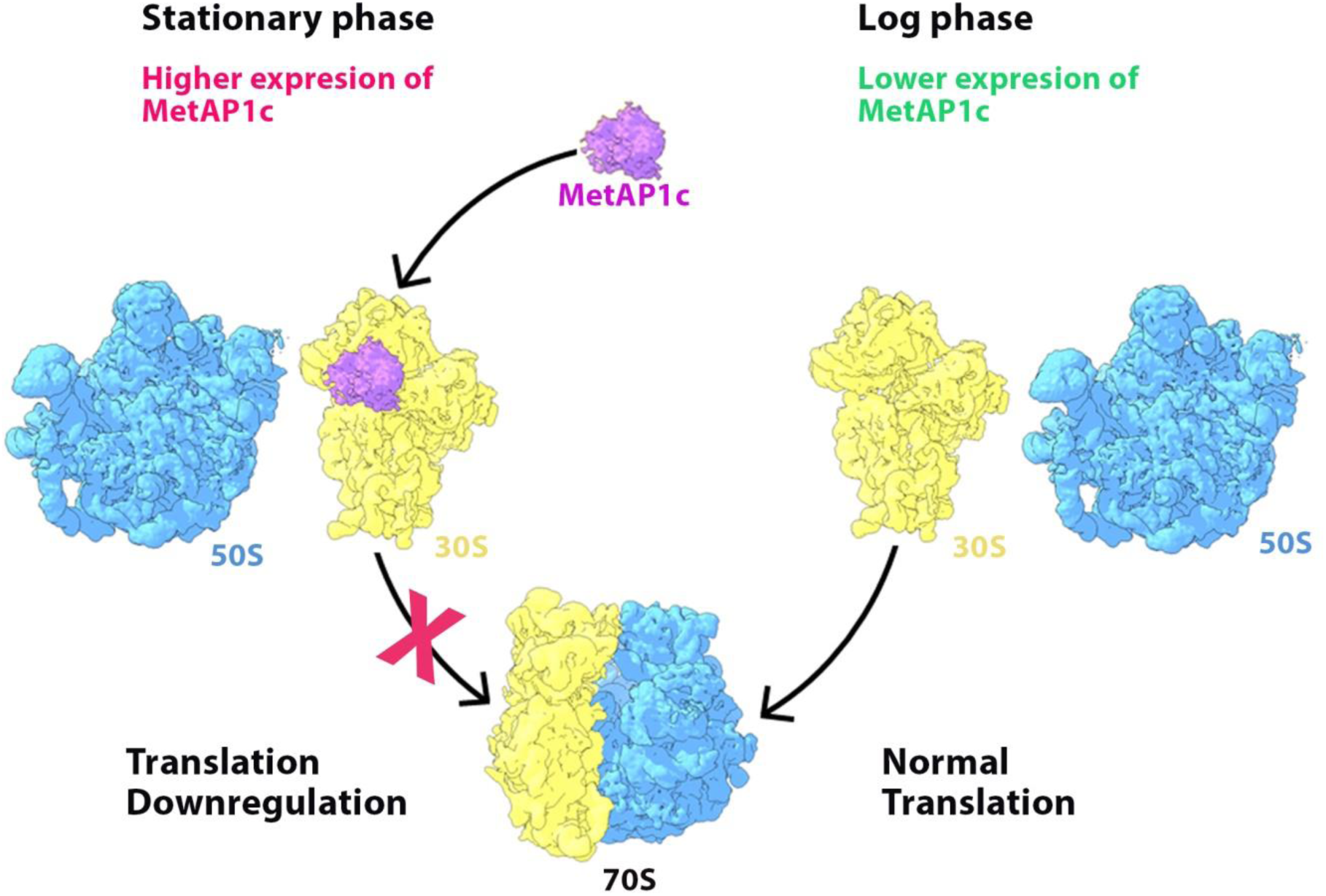
Schematic representation of the mechanism of MsmMetAP1c-mediated anti-association. During the stationary growth phase, the level of MsmMetAP1c is heightened, potentially leading to increased binding to the 30S subunit. However, this increased binding of MsmMetAP1c to the 30S subunit disrupts the formation of active 70S ribosomes by inhibiting its association with the 50S subunit, ultimately resulting in a reduced rate of translation. During the log phase of growth, the low level of MsmMetAP1c results in reduced binding to the 30S subunit. Consequently, this facilitates the smooth association of the 50S subunit with the 30S subunit, forming an active 70S ribosome essential for maintaining translation rates.

## Materials and Method

Unless mentioned otherwise, all reagents and kits were purchased from Sigma-Aldrich.

### PCR amplification and cloning of genes

The genes encoding mycobacterial Methionine aminopeptidase 1a (MetAP1a), Methionine aminopeptidase 1c (MetAP1c), Del40 MetAP1c and *E. coli* MetAP were PCR amplified using specific primers (Table) and sub-cloned into antibiotic resistant vectors containing requisite restriction sites. Chimeric MetAP1a was prepared by cloning it in two steps. The N-terminal extension sequence was first cloned into pET21c vector and MsmMetAP1a was then inserted into the same plasmid downstream of the N-terminal insert using the same primer sequence.

**Table M1.**
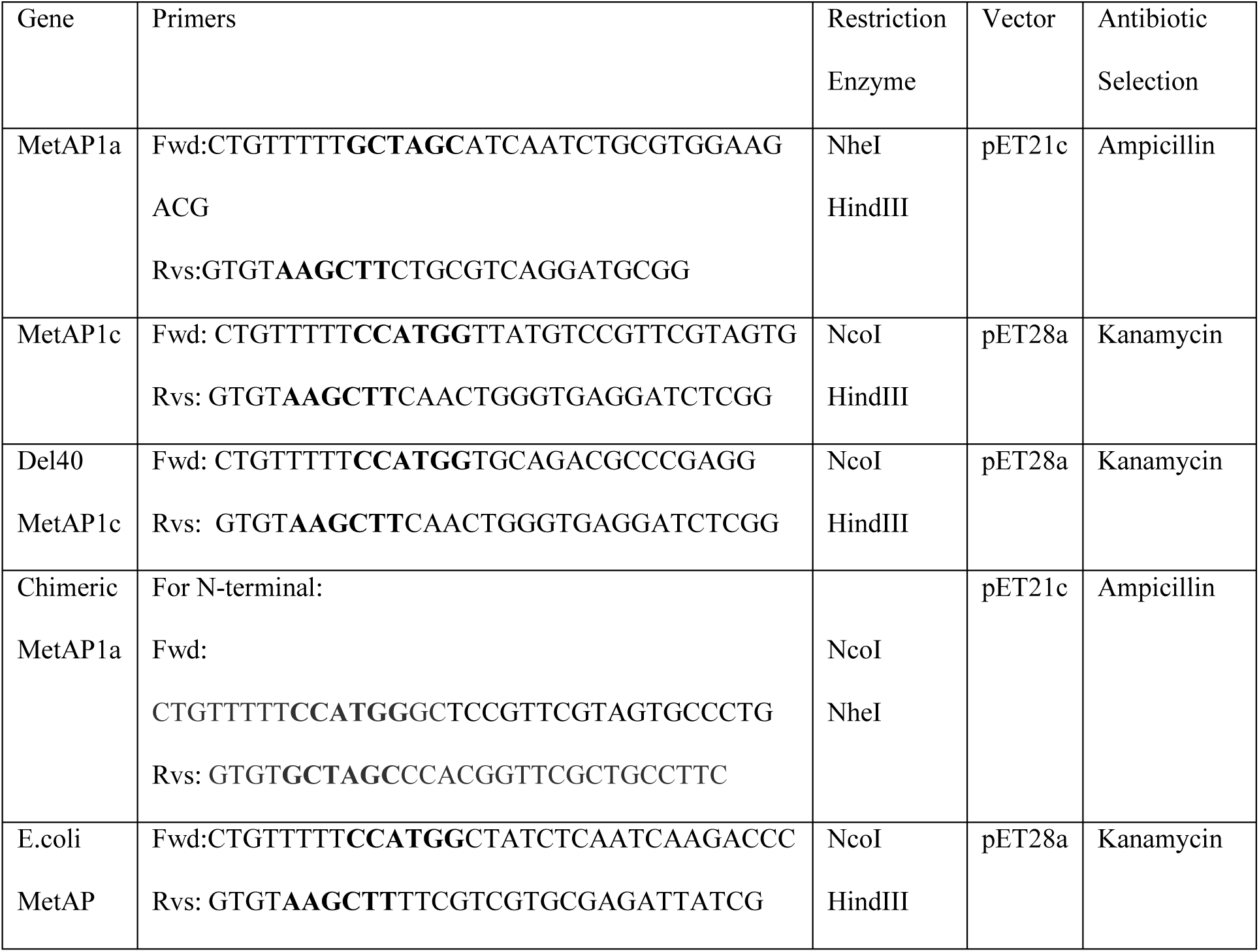
shows the primers and restriction enzymes used for cloning.

### Expression and purification of recombinant proteins

The positively cloned vectors were transformed into expression hosts and the protein expression was optimized by standardizing the IPTG dose and induction parameters (Table). Different buffers were used to purify the 6x His tagged proteins using Ni-NTA chromatography. Finally, the proteins were concentrated using Amicon Ultra-4 Centrifugal Units and stored at -80° C with 10% glycerol.

**Table M2.** shows expression host and induction parameters.

| Protein | His Tag | Expression Host | IPTG Dose and Temp |
| --- | --- | --- | --- |
| MetAP1a | C-Terminal | *E.coli* BL21 (DE3) | 0.1mM; 25°C for 16hrs |
| MetAP1c | C-Terminal | *E.coli* BL21 (DE3) | 0.5mM; 37°C for 2hrs |
| Del40 MetAP1c | C-Terminal | *E.coli* BL21 (DE3) | 0.5mM; 37°C for 2hrs |
| Chimeric MetAP1a | C-Terminal | *E.coli* BL21 (DE3) | 0.1mM; 16°C for 18hrs |
| *E.coli* MetAP | C-Terminal | *E.coli* BL21 (DE3) | 0.1mM; 25°C for 16hrs |

### Purification of Mycobacterial 70S, 50S and 30S ribosomes

Ribosome was purified based on our lab protocol with some modifications. Briefly, for 70S ribosome isolation, *Mycobacterium smegmatis* mc^2^155 cells were harvested from late log phase and washed thrice with Buffer A [20 mM Tris–HCl (pH 7.5), 10 mM MgOAc_2_, 100 mM NH_4_Cl, 5mM βME]. The pellet was again dissolved in buffer A with 1mg/ml lysozyme and lysed by sonication. The lysate was centrifuged to remove the debris and the crude ribosome was obtained by ultra-centrifugation using a swing out rotor Thermo Scientific™ AH-629 (154,000g, 4 °C, 2 h). The crude ribosome pellet was stored overnight in Buffer B [20 mM Tris–HCl (pH 7.5), 10 mM MgOAc_2_, 30mM NH_4_Cl, 5mM βME]. Pellet was re-suspended in Buffer B and homogenized for 1 h in the presence of 1M NH_4_Cl, after which the ribosomal preparation was centrifuged several times (15,000g, 4 °C, 20 min) to eliminate any visible pellet. Supernatant solution was ultra-centrifuged using swing-out rotor Thermo Scientific™ AH-629 (154,000g, 4 °C, 2 h). The pellet was collected and kept overnight in Buffer B and then re-suspended. Sample was loaded on top of a 10%–40% linear sucrose gradient in Buffer B and centrifuged using swing-out rotor Thermo Scientific™AH-629 (154,000g, 4 °C, 4hrs 30min). Gradient was collected from bottom to top and relevant fractions were pooled together. The 70S ribosome fraction was concentrated in Buffer B using 100 KD Amicon® Ultra-4 Centrifugal Filter Units (Merck Millipore) and stored at -80 °C.

For 50S and 30S isolation, the re-suspended pellet after 2^nd^ round ultra-centrifugation was dialyzed with Buffer B containing 1mM MgOAc_2_ for 8hrs and loaded on top of 10%-40% sucrose gradient. It was centrifuged using swing-out rotor Thermo Scientific™AH-629 (154,000g, 4°C, 9hrs 30min). The fractions were collected in a similar manner and relevant fractions were pooled together. The fractions were concentrated and stored.

### Gel electrophoresis and western blotting

Ribosome samples with or without bound proteins were loaded on a 15% denaturing SDS-PAGE gel and stained with Coomassie blue for visualizing protein bands. Equal amount of RNA was loaded in each lane by normalizing using OD260 values.

Western blot analysis was performed for ribosomes with or without the bound recombinant proteins. Blots were probed with MetAP1c and MetAP1a specific antibody (Bio Bharati Life Sciences) to detect MetAP1c and MetAP1a respectively. Del40 MetAP1c was detected with MetAP1c antibody, *E.coli* MetAP and Chimeric MetAP1a with MetAP1a specific antibody. Apart from normalizing by OD260, small ribosomal subunit protein S6 was used as a loading control (Cell Signaling Technology #2217). Binding of the secondary HRP-conjugated anti-rabbit antibody (Millipore, USA) was analyzed using ImmunoCruz (Santa Cruz Biotechnology Inc., USA).

### Phase-specific expression of MsmMetAP’s

To understand the phase specific protein expression levels, *M. smegmatis* cells were harvested at early log (OD600 value 0.6), late log (OD600 value 1.2) and stationary (OD600 value 2.5) phase. The cells were dissolved in Buffer B, lysed by sonication and 100ug of protein was loaded in each lane of SDS-PAGE. Since GAPDH, MsmMetAP1a and MsmMetAP1c have almost identical molecular weight, replica gels were prepared. One was probed with anti-GAPDH antibody (Invitrogen #MA5-15738), and the other with anti-MetAP1a or anti-MetAP1c antibody.

### Ribosome co-sedimentation assay

70S ribosome and ribosomal subunits were incubated with 25 folds excess recombinant protein in Buffer B. The sample was kept at 37°C for 15mins followed by 4°C for 15mins. It was then loaded on top of 10%-40% linear sucrose gradient and subjected to ultra-centrifugation for 4hr30mins. The fractions were collected, analyzed, pulled together and concentrated using 10 KD Amicon® Ultra-4 Centrifugal Filter Units (Merck Millipore). The complex and free ribosome were resolved in 15% SDS PAGE gel. The binding of proteins with ribosome was validated by Coomassie staining and Western Blot.

### Ribosomal subunit association assay

50S and 30S subunits were mixed at equimolar concentration and incubated in HMA-10 buffer (20mM HEPES pH7.6, 60mM NH_4_Cl, 10mM MgOAc_2,_ 5mM β-mercaptoethanol) at 37°C for 30mins. MsmMetAP1c-30S, MsmMetAP1a-30S, Del40MetAP1c-30S and Chimeric MetAP1a-30S pre-incubated complexes were also incubated with equimolar 50S subunit under similar condition. The samples were then loaded on top of 10%-40% linear sucrose gradient and run on a swing-out rotor Thermo Scientific™AH-629 (154,000g, 4 °C, 5hrs 30min). The fractions were collected and analyzed to obtain the ribosomal profile.

### Light scattering assay

60nM of 30S subunit was mixed with 900nM of MsmMetAP1a or MsmMetAP1c in HMA-10 buffer. Free 30S ribosomal subunit was used as a control.60nM of 50S subunit was added to both the sets and light scattering was measured at 350nm for 5mins in Hitachi F-7000 Fluorescence Spectrophotometer.

### *In-vivo* ribosomal binding studies

pLam12 vector containing MsmMetAP1a and MsmMetAP1c gene was transformed into *M. smegmatis* mc^2^155 cells by electroporation (37). The positively transformed cells were selected using kanamycin and induced with 0.1% acetamide at OD600 value of 0.8. After 3hrs of induction, the cells were harvested and ribosome was isolated in HMA-10 buffer using a sucrose density gradient based approach. The 70S, 50S and 30S fractions were separated, concentrated and run on SDS-PAGE to analyze the bound protein level.

### Luciferase assay

The S30 T7 High-Yield Protein Expression System of Promega was used for this assay. The reaction mixture was prepared as per protocol provided by the manufacturer. 500ng of recombinant plasmid containing luciferase gene was added in each reaction set. Varying concentration of MsmMetAP1a, MsmMetAP1c, Del40 MetAP1c and Chimeric MetAP1a were added to the reaction mixture and finally the amount of luciferase synthesized was analyzed by measuring firefly luminescence using luminometer (Promega).

### Statistical analysis

All graphs and statistical analysis were generated using Prism (v5.00) software (GraphPad, San Diego, CA). Unpaired *t* tests were used for analysis. P values of <0.05 were considered statistically significant. Error bars indicate means ± standard deviation (SDs).

### MsmMetAP1a model generation

The model of *Mycobacterium smegmatis* MetAP1a was generated using Phyre2 (38).

### Cryo-EM sample preparation and data collection

*M. smegmatis* 30S-MetAP1c complex was prepared by adding 25 folds excess protein to 30S subunit in HMA-10 buffer. After 15mins incubation at 37°C, the reaction mixture was cooled to 4°C and subjected to ultracentrifugation for 5hrs30mins. 30S fractions bound to MsmMetAP1c were pulled and used for cryo-EM grid preparation. Gold grids were used for data collection. The high-resolution data was collected at National Electron Cryo-Microscopy Facility, Bangalore Life Science Cluster (BLiSc), Bangalore using 300 keV Titan Krios (Thermo Fischer) with Falcon 3 direct electron detector camera. 4545 movie stacks were collected with 20 movie frames per stack at 1.07 Å/pixel with an electron dose of 1.035 e/Å^2^/frame.

### Cryo-EM data processing

Cryo-EM data processing was done using Relion-3.1.0 (39). Motion correction was done using Relion’s own implementation MotionCor2 (40), followed by CTF Estimation using CCTFFIND4 (41). Autopicked particles were subjected to iterative 2D classification and the best 2D classes were selected, corresponding to roughly 1,16,000 30S particles. These particles were subjected to 3D classification and classes which displayed proper 30S head were selected and pooled. Following further 2D classification, 3D classification, CTF refinement and particle polishing, a final particle set of 56,923 MsmMetAP1c bound 30S particles were refined to a resolution of 5Å (FSC criteria 0.143) (42).

### Model building and structure analysis

The PDB structure of *M. smegmatis* 30S subunit (PDB: 5ZEU, (19)) was docked into the map using Chimera (43). The MsmMetAP1c model generated using AlphaFold2 (44) software was used for fitting into the protein density. The model was built using WinCoot (version 0.9.8.1) (45) and refined using Phenix Real Space Refinement (46) iteratively. Model validation was done using MolProbity (47) in PHENIX. Figures were prepared using Chimera (43), ChimeraX (48), PyMOL (Schrödinger, LLC.) and WinCoot.

## Data Availability

The atomic coordinate has been deposited in the Protein Data Bank under the accession code 8YP6. The respective cryo-EM volume has been deposited in the Electron Microscopy Data Bank under the accession code EMD-39462.

## Acknowledgements

This work was primarily supported by SERB, DST (India) sponsored projects (CRG/2019/001788) and (SPF/2021/000141), CSIR Niche Creating Project (NCP) MLP-139 and CSIR-Indian Institute of Chemical Biology (CSIR-IICB), Kolkata, India. We greatly acknowledge the National Electron Cryo Microscopy facility at the Bangalore Life Sciences Cluster (DBT/PR12422/MED/31/287/2014), NCBS, Bangalore, DBT (grant no.BT/PR15017/BRB/10/1445) and the Central Instrumentation Facility (CIF) of CSIR-IICB. We sincerely thank Dr. Sucharita Bose and Dr.Vinothkumar Kutti Ragunath for helping in grid preparation and high resolution single particle data collection. AB and KS acknowledge DBT and CSIR, India, respectively for research fellowship.

## Author contributions

JS conceived the project. AB designed the experiments in consultation with JS, carried out all the biochemical experiments and prepared the illustrations. AB and KS did the cryo-EM 3D image processing. JS, AB and KS analysed the data and wrote the paper.

