## Supplemental Data for "Mycobacterial methionine aminopeptidase type 1c moonlights as an anti-association factor on the 30S ribosomal subunit"

[3]Corresponding author

**Table S1:**

| Ribosome Complex | Msm MetAP1c-30S |
|---|---|
| **Database submission** | |
| EMDB ID | EMD-39462 |
| PDB ID | 8YP6 |
| **Data Collection** | |
| Microscope | Titan Krios |
| Camera | Falcon III (4k x 4k) |
| Nominal Magnification | 75000x |
| Voltage (keV) | 300 |
| Electron Dose (e/ Å$^2$) | 20.7 |
| Defocus Range (um) | -1.8 to -3.3 |
| Pixel Size (Å) | 1.07 |
| **Cryo-EM Reconstruction** | |
| Refined Map Resolution (Å) | 5Å |
| Final Particle Number | 56,923 |
| Symmetry | C1 |
| FSC Threshold | 0.143 |
| Resolution Metric | FSC |
| **Model Composition** | |
| Chains | 20 |
| Non-hydrogen Atoms | 51568 |
| Protein Residues | 2448 |
| Nucleotide Residues | 1506 |
| Ligands | 0 |
| **Root-mean-square Deviations** | |
| Bond Length (Å) | 0.004 |
| Bond Angle (°) | 0.785 |
| **Validation** | |
| MolProbity Score | 2.52 |
| Clash Score | 33.28 |
| Rotamer Outliers (%) | 0.84 |
| **Ramachandran Plot** | |
| Outliers (%) | 0.46 |
| Allowed (%) | 8.42 |
| Favoured (%) | 91.12 |

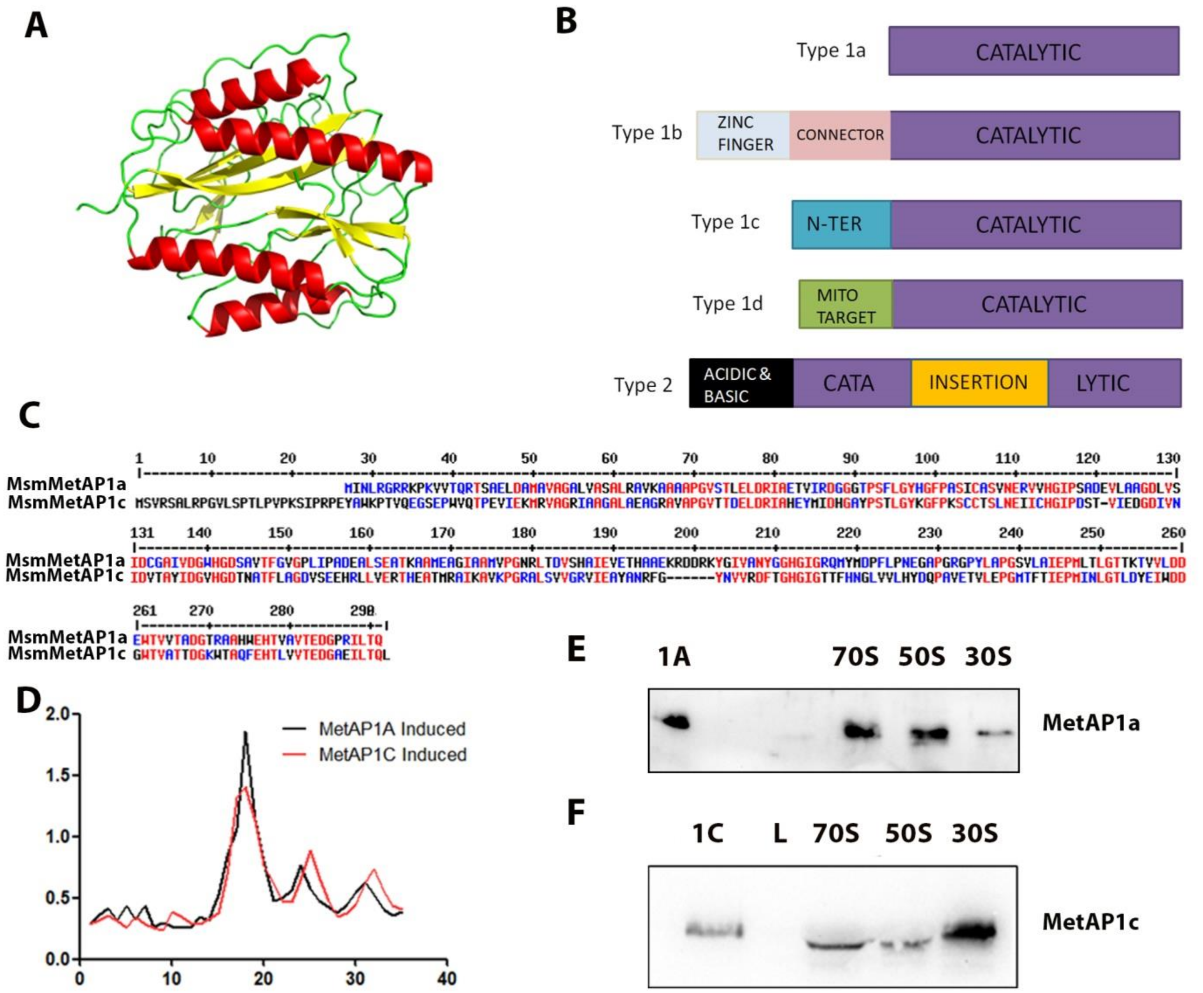


**Figure S1. Different Methionine aminopeptidases (MetAPs), structure and function**
The conserved "pita-bread" fold of methionine aminopeptidase is depicted in (A). A schematic representation of the different types of MetAPs highlights variations in the N-terminal and catalytic core (B). Sequence alignment between MsmMetAP1a and MsmMetAP1c reveals the presence of the N-terminal extension in MsmMetAP1c (C). In *E. coli* cells containing recombinant plasmid encoding mycobacterial MetAP1a and MetAP1c, induction was performed, followed by isolation of ribosomes. The resulting ribosomal profile (D) suggests a decrease in the 70S population upon MsmMetAP1c induction. Further analysis of the ribosomal fractions reveals the binding preference of MsmMetAP1a towards the 70S ribosome and 50S subunit (E), whereas MsmMetAP1c exhibits preference towards the 30S subunit (F).
(In the ribosomal profile of panel D, X axis corresponds to fraction number and Y axis corresponds to OD260)

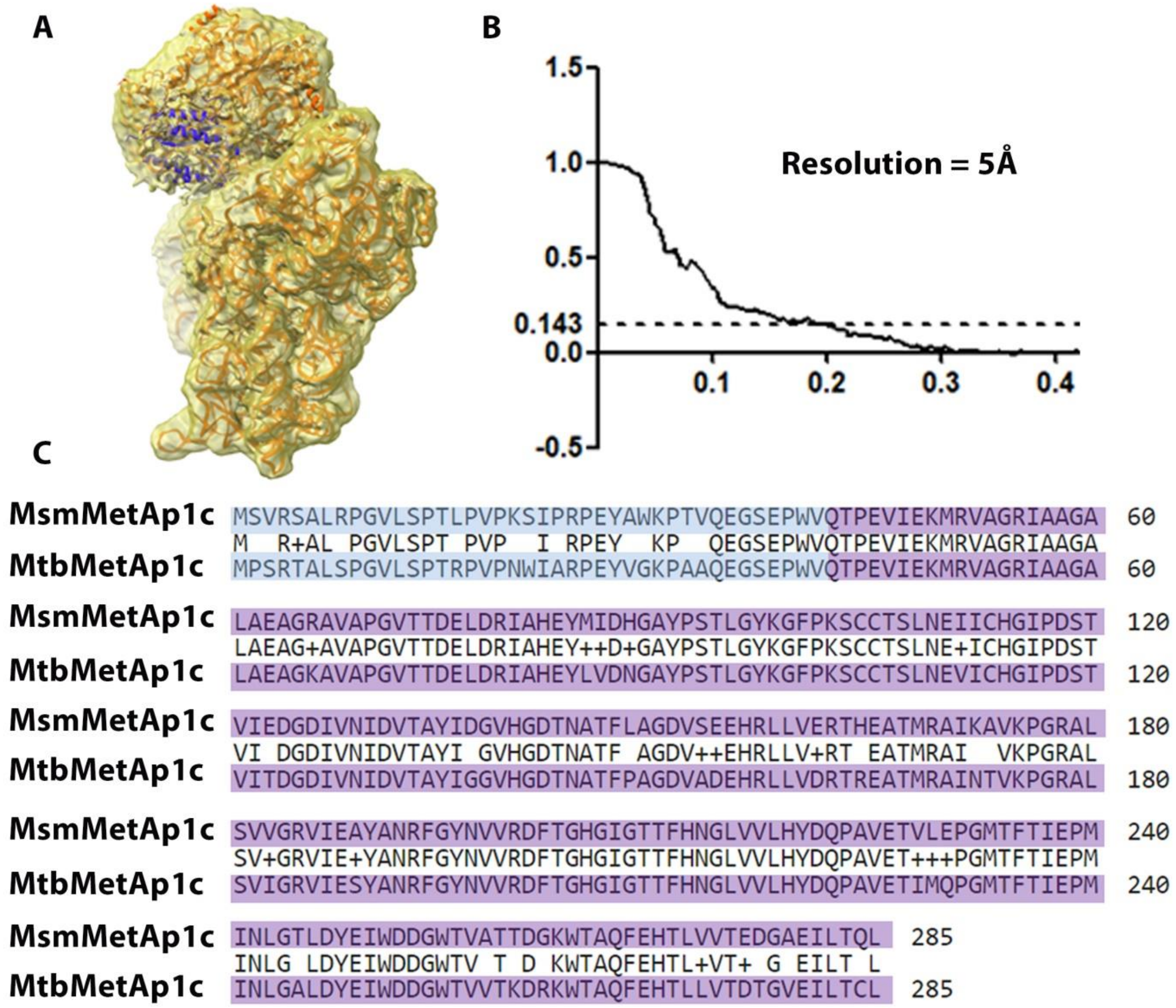


**Figure S2. Cryo-EM reconstruction of MsmMetAP1c-30S subunit complex and sequence alignment**

The map model overlay displays the overall fitting of the complex (A). The Fourier Shell Correlation (FSC) plot demonstrates the resolution of the obtained map as 5Å (B). The sequence alignment of *M. smegmatis* MetAP1c and *M. tuberculosis* MetAP1c underscores the overall similarity between the proteins (C).

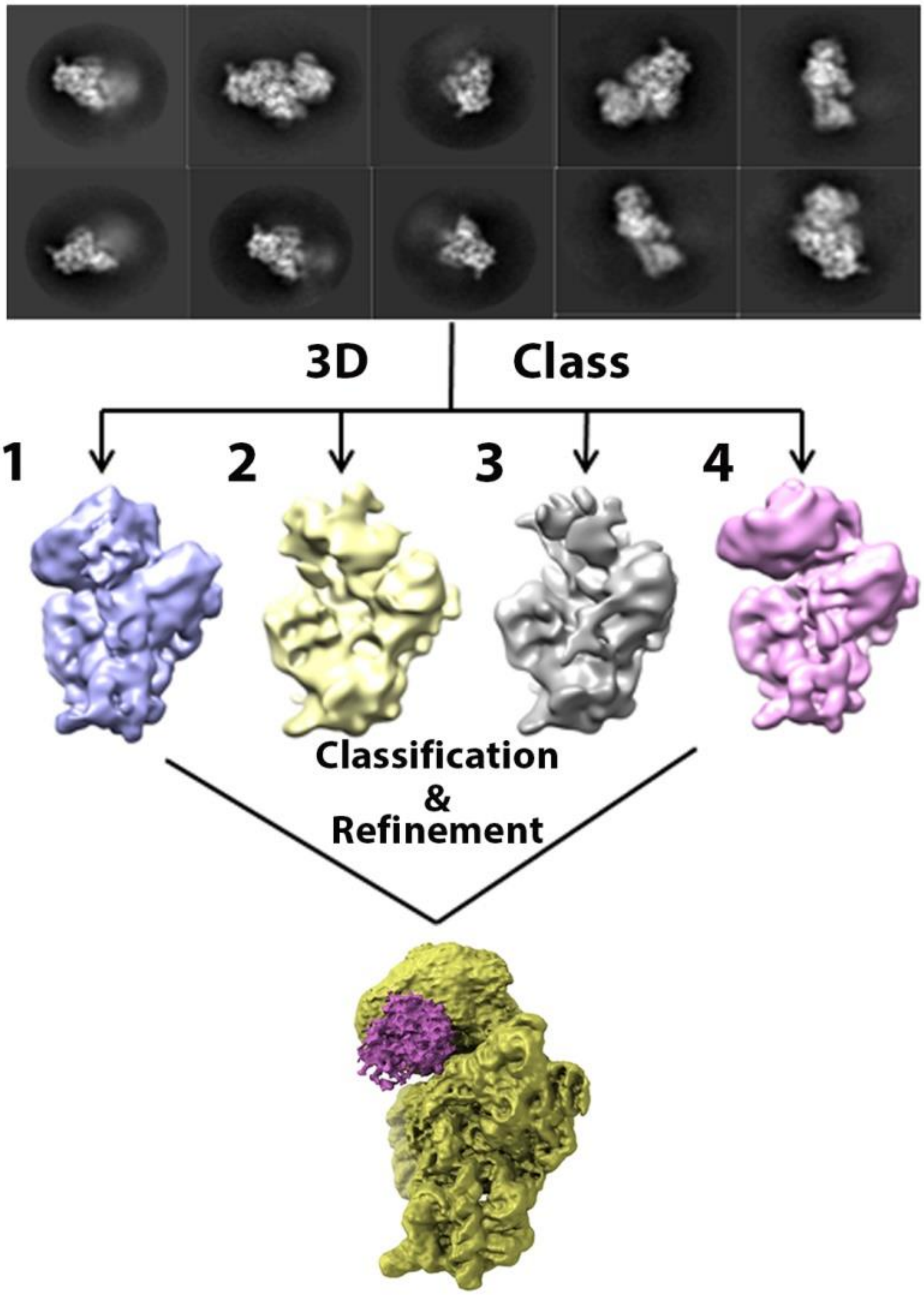


**Figure S3. 2D and 3D classifications of MsmMetAP1c-30S subunit dataset**
The 2D classification reveals distinct classes, some displaying proper 30S subunit heads while others exhibit hazy heads. Subsequent 3D classification segregates particles with distorted heads (classes 2 and 3) from those with proper heads (classes 1 and 4). Particles from classes 1 and 4 are grouped together, subjected to further classification and refinement, ultimately yielding the final MsmMetAP1c bound 30S map.

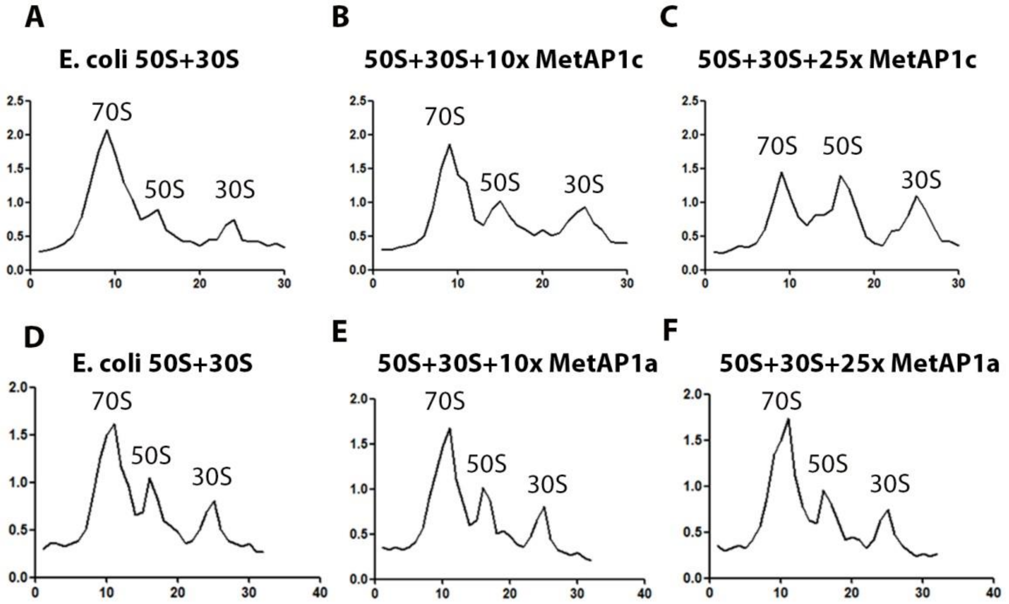


**Figure S4. Association profiles of *E. coli* ribosomal subunits**

*E. coli* 50S and 30S subunits associate to form the 70S ribosome (A). This process is hindered by the presence of MsmMetAP1c. When 10-fold (B) and 25-fold (C) excess MsmMetAP1c is introduced, a noticeable reduction in 70S ribosome formation is observed. In contrast, the presence of MsmMetAP1a does not affect the efficiency of association. In the control set (D), 70S ribosome formation proceeds normally. However, the addition of 10-fold excess MsmMetAP1a (E), and even with 25-fold excess MsmMetAP1a (F), does not lead to any significant reduction in 70S ribosome formation compared to the control condition.
(In the association profiles, X axis corresponds to fraction number and Y axis corresponds to OD260)

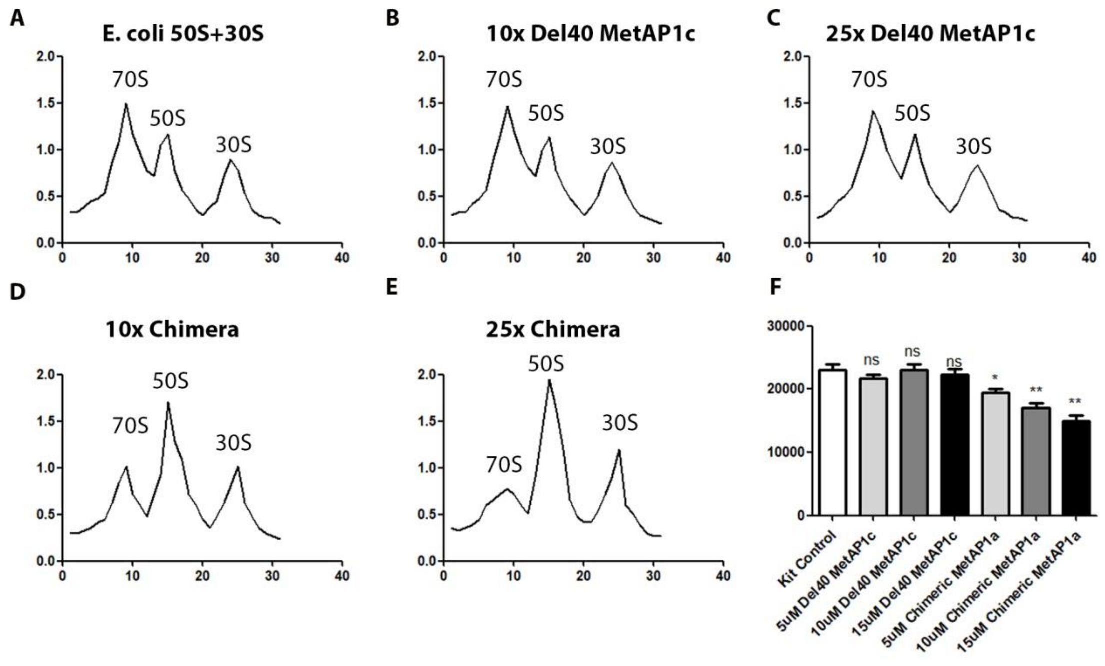


**Figure S5. Anti-association property of Del40 MetAP1c and Chimeric MetAP1a**
Del40 MetAP1c lacks the anti-association property. *E. coli* 50S and 30S subunits associate to form the 70S ribosome (A). Comparison of this profile with 10-fold (B) and 25-fold (C) excess Del40 MetAP1c treated sets reveals no change in pattern. In contrast, 10-fold (D) and 25-fold (E) excess Chimeric MetAP1a leads to diminished 70S formation. This reduction is further supported by the decrease in luciferase synthesis. As the amount of Chimeric MetAP1a increases, there is a corresponding decrease in firefly luminescence, indicating lower 70S ribosome formation (F).
(In the association profiles, X axis corresponds to fraction number and Y axis corresponds to OD260)

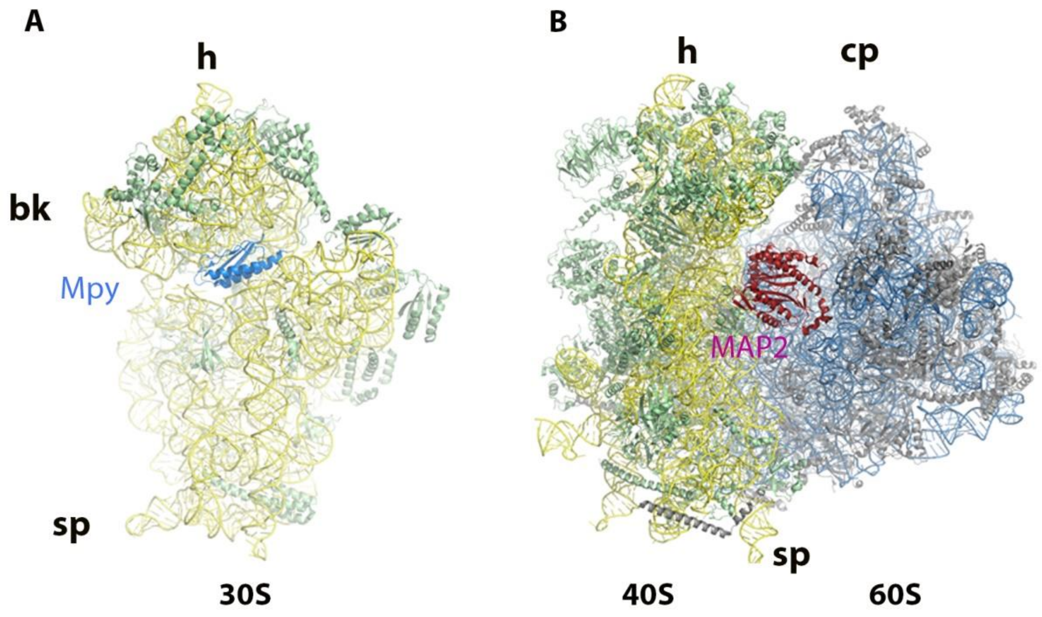


**Figure S6. The interactions of Mpy and MAP2 with 30S subunit and 80S ribosome respectively**

The binding of Mpy (blue) with 30S subunit (16S rRNA in yellow, proteins in green) does not alter the conformation that will result in hindrance to 70S formation (A). MAP2 (red) binds to 80S ribosome by interacting with small ribosomal protein S12. The presence of MAP2 does not lead to a steric clash with the 60S subunit thus has no effect in 80S formation (B).

h: Head; bk: Beak; sp: Spur; cp: Central Protuberance
